# Predicting T cell receptor antigen specificity from structural features derived from homology models of receptor-peptide-major histocompatibility complexes

**DOI:** 10.1101/2021.05.19.444843

**Authors:** Martina Milighetti, John Shawe-Taylor, Benjamin M Chain

**Affiliations:** Division of Infection and Immunity, University College London, London, United Kingdom; Cancer Institute, University College London, London, United Kingdom; Department of Computer Science, University College London, London, United Kingdom

## Abstract

The physical interaction between the T cell receptor (TCR) and its cognate antigen causes T cells to activate and participate in the immune response. Understanding this physical interaction is important in predicting TCR binding to a target epitope, as well as potential cross-reactivity. Here, we propose a way of collecting informative features of the binding interface from homology models of T cell receptor-peptide-major histocompatibility complex (TCR-pMHC) complexes. The information collected from these structures is sufficient to discriminate binding from non-binding TCR-pMHC pairs in multiple independent datasets. The classifier is limited by the number of crystal structures available for the homology modelling and by the size of the training set. However, the classifier shows comparable performance to sequence-based classifiers requiring much larger training sets.

## 2 Introduction

T cells are key players of adaptive immunity. They are activated by the recognition of a cognate peptide, a short stretch of amino acids which is displayed on a major histocompatibility complex molecule (MHC, pMHC when bound to peptide). The recognition occurs via the T cell receptor (TCR), which is composed of two chains (normally an *α* and a *β*), both of which are generated by a process of random recombination and selection. The recombination gives rise to 3 hypervariable regions, the complementarity-determining regions - CDR1, CDR2 and CDR3. Among the three regions, CDR3 is the most variable as it is found at the junction of V(D)J recombination, and it can therefore incorporate a number of non-template insertion and deletion events, whilst CDR1 and CDR2 depend on the V gene selected in the recombination process and have therefore a lower number of possible sequences.

A number of TCR-pMHC complexes have been crystallised and the structures solved and they are collected in the Structural T-Cell Receptor Database (STCRDab, Leem et al. 2018). They have given us deeper understanding of TCR-pMHC interactions and how these are impacted by mutations, but also how structure and function are related. Examples include how cross-reactivity between bacterial and self antigens can drive disease (Petersen et al. 2020), how binding mode can give different specificity profiles to TCRs binding the same peptide (Coles et al. 2020), and how binding orientation is determined by how the peptide is presented by the MHC (Singh et al. 2020).

The existing structures can also be mined for information on how the TCR interacts with the pMHC complex. By looking at the TCR residues that fall within 5Å of the peptide in a number of published TCR-pMHC structures, both Glanville et al. 2017 and Ostmeyer et al. 2019 showed that the CDR3 is the region that makes the most extensive contacts with the peptide. These regions of contact are normally short stretches of 3 or 4 consecutive amino acids within the CDR3. Moreover, they noted that whilst the TCR*β* always made contacts, there are multiple instances were the TCR*α* is not within contact distance of the peptide. It has also been shown that TCRs which recognise the same peptide share motifs and sequence characteristics in the CDR3 (Thomas et al. 2014; Cinelli et al. 2017; Glanville et al. 2017; Dash et al. 2017).

The ensemble of TCRs that are present within an individual at any point in time is called the TCR repertoire. Different sequences are found at widely different frequencies, ranging from a few hundred copies to 10^9^ copies for the larger T cell clones, which make up up to 1% of the total repertoire. Differences in clone size can arise both in the naive repertoire, by convergent recombination (whereby an amino acid sequence is likely to be produced by the process of recombination - normally with short CDR3 and few nucleotide insertions, Venturi et al. 2006; Britanova et al. 2014) or because of the power-law distribution of naive T cell clones produced by the thymus (Greef et al. 2020); or in the memory repertoire by convergent selection, whereby similar sequences are expanded because they are responding to the same antigen, Pogorelyy et al. 2018). Greef et al. 2020 estimates the maximum effect of generation probability to be around 10^7^, which is two order of magnitudes smaller than the largest observed clone sizes, suggesting a role for expansion during the immune response. By focusing solely on the CDR3, it can be shown that during an immune response, expanded TCR clones are frequently part of clusters of sequences that are more similar to each other than might be expected by random sampling of the repertoire (Joshi et al. 2019; Pogorelyy et al. 2019; Marcou et al. 2018).

This observation of antigen-driven TCR sequence clustering has been used to build algorithms such as GLIPH (Glanville et al. 2017) and TCRdist (Dash et al. 2017), which can build sequence motifs starting from a cluster of TCRs known to recognise the same peptide and which are then able to find other TCRs responding to the same peptide. More recently, Tong et al. 2020 have shown that sequence information encoded in the form of overlapping amino acid quadruplets can be used to create a multi-class prediction algorithm able to correctly assign TCR-pMHC pairs.

In the same way that conserved sequence motifs characterise TCRs recognising the same antigen, we hypothesise that there will be structural features of the TCR/antigen interface which are conserved in the interactions. Such conserved structural features could be leveraged to gain a better understanding of the TCR-pMHC interaction and to reca-pitulate and improve what has been learnt from looking purely at sequence information. Our understanding of the physical interactions between TCRs and pMHC is, however, limited to the set of solved and published crystal structures. The STCRDab currently reports about 400 entries for *αβ* TCR-pMHC complexes, and 120 different peptides, which is clearly a tiny subset of all the possible TCR-pMHC interactions that can exist. To solve this problem, a number of tools have been developed and subsequently optimised to predict the structure of a TCR-pMHC complex based on its sequence. One of these is TCRpMHCmodels (Jensen et al. 2019), which exists as a free online user interface. TCRpMHCmodels leverages LYRA (Klausen et al. 2015) to model the TCR structure and MODELLER (Fiser and Šali 2003) to predict the pMHC structure, to then combine them together by using a third set of templates for the TCR-pMHC complex overall. Tools like TCRpMHCmodels, although still limited by the amount of information that has been published, allow us to delve deeper into the structural relationships between the TCR and the pMHC.

We show here that a combination of structural and sequence features can be incorporated into a machine learning algorithm to discriminate binding and non-binding TCR-pMHC pairs. The classifier presented is limited by the performance of the homology modelling, but, unlike any of the previous work reviewed above, it does not rely on the identification of a set of TCRs binding to a specific peptide to be able to predict whether other TCRs will bind to that same peptide, but rather learns some general rules which can predict TCR interaction with completely novel peptides.

## 3 Methods

### 3.1 Datasets

The available crystal structures for TCR-pMHC complexes were retrieved from STCRDab (http://opig.stats.ox.ac.uk/webapps/stcrdab/, Leem et al. 2018). The dataset (referred to as STCRDab or PDB set) was refined to include only one complex per crystal, remove *γδ* TCRs and remove non-peptide antigens. The set was then checked for repeat sequences. For the classifier step, TCRs binding MHC class II complexes were removed as these cannot be modelled by TCRpMHCmodels. To create non-binding TCR-pMHC pairs, random TCR-pMHC pairs were created from the available pool, under the condition that the pMHC from the random pairing was not the same as the original one.

The 10XGenomics dataset was downloaded from the 10XGenomics website (CD8+ T cells of Healthy Donor 1, *A New Way of Exploring Immunity - Linking Highly Multiplexed Antigen Recognition to Immune Repertoire and Phenotype.*). For each TCR, binding (or absence of binding) to an epitope was defined as in the Application Note provided by 10X Genomics. Briefly, a specific binding event was defined as having UMI count higher than 10 and greater than 5 times the highest negative control for that TCR clone. When a TCR clone was assigned multiple barcodes, the UMI counts for each tetramer were summed to determine overall binding. If these conditions were true for more than one peptide, the TCR was called a binder for each of the epitopes.

The Dash dataset (generated by Dash et al. 2017) was obtained from the VDJDb dataset. Duplicate TCR-pMHC pairs were removed. Each unique TCR clone was paired with each pMHC in the dataset, making 1 binding and 9 non-binding complexes per TCR.

The set of experimental constructs (expt) consists of a set of experimentally-validated peptide-specific TCR constructs with cognate peptide, which have been characterised functionally: 2 CMV-reactive TCRs (NLVPMVATV peptide), 3 influenza-reactive TCRs (2 HA1-reactive - peptide VLHDDLLEA - and 1 HA2-reactive - YIGEVLVSV peptide), 1 EBV-reactive TCR (peptide CLGGLLTMV) from Thomas et al. 2019 and Chatterjee et al. 2019; A7 TCR and 3 affinity-matured TCRs from A7 which recognise pTax as well as pHud peptides (LLFGYPVYV and LGYGFVNYI, respectively) (Thomas et al. 2011); two TCRs identified as neoantigen-reactive in Joshi et al. 2019 and two mutated versions of these, which have been shown not to bind the neoantigen (unpublished data, A. Woolston, personal communication, 2020). To create the non-binders, each TCRs was matched with each pMHC in the pool, as well as with peptide WT235 (control peptide in Thomas et al. 2019, CMTWNQMNL) and peptide WTlung (FAFQEDDSF, wild-type peptide for the neo-antigen McGranahan et al. 2016).

A dataset of TCR-pMHC complexes with experimentally-determined affinity was retrieved from the ATLAS (http://atlas.wenglab.org/web/index.php, Borrman et al. 2017) to evaluate the impact of affinity on the classifier performance. Any TCR-pMHC pair with undetectable binding (*K_d_* labelled as *n.d.*) was called a non-binder whilst all other complexes were labelled binders regardless of the detected *K_d_*.

Finally, a dataset of TCR-pMHC complexes with epitopes that are neither present in our training set nor in the training set of the tools we benchmarked against was downloaded from the latest version of the VDJDb (Bagaev et al. 2020). As for the PDB set, negatives were created by shuffling of TCR-pMHC pairs in the set.

### 3.2 Homology modelling and feature extraction

Each structure (both binders and non-binders) in these datasets was homology-modelled with TCRpMHCmodels (which was kindly provided in command-line form by the authors, Jensen et al. 2019) in its default settings and submitted to the feature-extraction pipeline.

To make the structures comparable, they were renumbered to the standardised IMGT numbering (Lefranc 1997) using ANARCI (Dunbar and Deane 2016). Moreover, the peptide residues were renumbered to 1-20, so that the central residues would be residues 10-11 in each complex.

For each TCR-pMHC, 5 sets of features were extracted, namely:

- minimum pairwise distances between each CDR residue and each peptide residue were calculated using BioPDB (Hamelryck and Manderick 2003). These capture the binding mode of the TCR-pMHC complex;
- energetic profile of pairwise CDR-peptide residues interactions was calculated using PyRosetta v2020.28+ (Chaudhury et al. 2010). The Rosetta energy function for context-independent residue-residue interactions was used to extract the following terms (scorefunction: talaris2014) from a PDB file from which the MHC complex was removed: attractive and repulsive van der Waals (atr, rep), electrostatic interactions (elec) and solvation energy (sol) (Alford et al. 2017). These are a representation of binding energy of the complex.
- Atchley factors (Atchley et al. 2005) were used to encode the sequences of the peptide and CDRs for each TCR-pMHC pair.

To evaluate the effect of homology modelling performance on the classifier presented, the structures were categorised as having or not having good homology modelling templates. This was defined based on the sequence homology to the most similar peptide template (*>* 45% sequence similarity to the best pMHC model template) and complex template (*>* 60% sequence similarity to the best complex template). These thresholds were chosen based on the results presented by Jensen et al. 2019.

To be noted that not all structures could be successfully modelled by TCRpMHC- models, and so we could not submit them to the feature extraction pipeline.

### 3.3 Multiple kernel learning

Each feature set was pre-processed separately. Missing values were imputed with the median value of the feature across the train set. Each feature was then scaled to have a value between 0 and 1 (sci-kit learn Minmax scaler, Pedregosa et al. 2011) and normalised. To properly represent and integrate the different features extracted from the structures, kernels were created separately for each subset of features. Moreover, instead of optimising a single kernel for each feature set, 7 Gaussian (rbf) kernels were created and combined, letting the MKL algorithm decide the weights for each kernel, as in Lauriola et al. 2017. The *γ* parameters for the 7 Gaussian kernels for each feature set were found as follows:

1. calculate the distance between all positive (binding, n) and negative (non-binding, m) examples in the train set

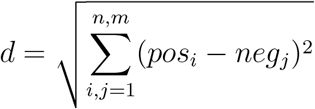
2. find *σ* values corresponding to 1^st^, 2^nd^, 5^th^, 50^th^, 55^th^, 98^th^ and 99^th^ percentile of distances
3. for each *σ*, calculate the *γ* as:

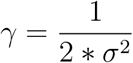

The kernels generated were combined by the EasyMKL algorithm as implemented in MKLPy to find an optimal combination (Aiolli and Donini 2015; Lauriola et al. 2017; Lauriola and Aiolli 2020), setting sci-kit’s learn SVC algorithm as a learner (Pedregosa et al. 2011). The *λ* parameter for EasyMKL was fixed to 0 and the optimal C parameter for SVC was searched in the range between 10*^−^*^5^ and 10^2^ by 10-fold (internal) cross-validation (CV) on the train set. This process was used both when a single feature set was evaluated (by combining the 7 kernels for the set) and when combining multiple feature sets (7 kernels for each set).

To estimate performance by cross-validation, the train set was split 70-30. 70% was used to optimise the model parameters by maximising the ROC AUC score and the remaining 30% was used for prediction. The procedure was repeated 10 times with different subsets of samples.

Out-of-sample performance was evaluated in the datasets outlined in section 3.1, by training the classifier on the whole of the training set.

### 3.4 Benchmarking against other classifiers

To evaluate the performance of the presented classifier compared to published classifiers in the field, we compared performance with ERGO (Springer et al. 2020) and ImRex (Moris et al. 2020) on the same validation sets. ERGO is available as a web tool (http://tcr.cs.biu.ac.il/), and the models trained on the VDJdb (Bagaev et al. 2020) were used for the benchmarking. ImRex is available as a GitHub repository (https://github.com/pmoris/ImRex), and the available model trained on the VDJdb was used for the predictions.

### 3.5 Data availability

The complete set of sequences used, as well as prediction results are provided as supplementary files.

## 4 Results

### 4.1 Extracting physical features from available TCR-pMHC complex structures allows interrogation of binding mode

We first established a systematic pipeline to extract structural information about the TCR-peptide interface from a dataset of solved structures downloaded from the Structural T Cell Receptor Database (Leem et al. 2018). The minimum pairwise distances between TCR and peptide residues, and their corresponding attractive and repulsive van der Waals forces (atr, rep), electrostatic interactions (elec) and solvation energies (sol) were estimated for each peptide-TCR complex as described in the methods.

Each feature extraction process yielded a matrix with information about pairwise contacts between residues in the TCR and residues in the peptide (Figure 1a). The distance fingerprints are easy to compare between different structures and can give insight into the binding mode for the complex: for instance, complexes 1AO7 (Garboczi et al. 1996) and 1MI5 (Kjer-Nielsen et al. 2003) (both MHC Class I) bind closer to the N terminus of the peptide, whilst 1D9K (Reinherz et al. 1999) has the TCR bound more centrally, and this is particularly evident in the *α* chain (Figure 1a and b).

**Figure 1:**
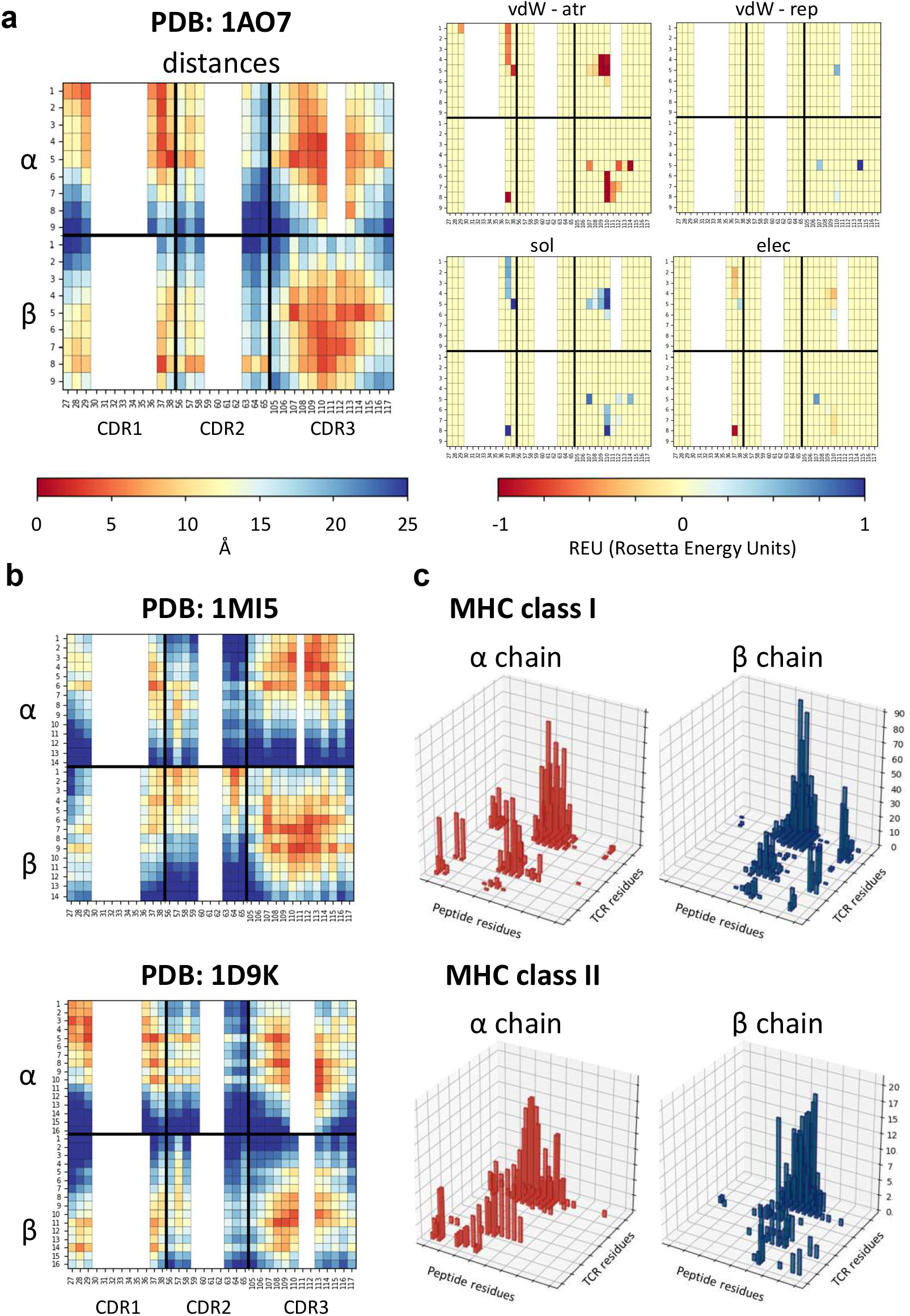
Feature extraction from PDB structures. **a.** Heatmaps showing the physical features extracted for structure 1AO7. In each heatmap, the top half refers to the *α* chain and the bottom half to the *β* chain. Each column is a CDR residue, each row is a peptide antigen residue and the colour of each square represents the value extracted for the CDR-peptide residue pair (i.e. top left-hand square of the distance panel is the distance between residue 1 on the peptide and residue 27 of the TCR*α* chain). Similar plots are shown for each energy term extracted - van der Waals attractive, van der Waals repulsive, solvent and electrostatic. **b.** Two other examples of distance fingerprints, a class I and a class II complex - 1MI5 (class I complex, EBV peptide) and 1D9K (class II complex, conalbumin peptide) - for comparison with 1AO7. Same scale as in a. **c.** Histograms showing the number of structures making a contact (less than 6Å) for each peptide residue-CDR residue pair, for alpha and beta chains separately, showed for class I and class II complexes. Peptide residues renumbered 1-20 for consistency as described in methods

We wondered whether any trends could be detected more generally and used the minimum pairwise distances to identify the distribution of interactions between TCR CDR residues and the peptide in class I and class II complexes (Figure 1c). While it is clear that interactions between TCR chains and antigen peptide are not confined to a single hotspot, some general patterns emerge. The TCR*α* chain, for example, tends to bind the N-terminus of the peptide, whilst the *β* binds towards the C-terminus, as has been reported previously (Garcia et al. 2009). Interestingly, while contacts were dominated by the CDR3 region of the TCR, we also detected contacts between CDR1 and CDR2 and peptide residues in a significant proportion of complexes. Moreover, more of the class I structures make contacts with the C-terminus of the peptide than class II. A similar pattern is also detected when looking at the energetic interactions (Supplementary Figure S1).

In order to look in more detail for potential conserved patterns with which to characterise the TCR-peptide binding surface, we calculated a PCA for each of the feature sets (distances and energy vectors) for all complexes (Figure 2a and Supplementary Figure S2a). The first dimension of the PCA of the minimum pairwise distances clearly identified the few examples where the TCR is in an inverse orientation relative to the peptide (stars, PDB: 4Y19 and 4Y1A Beringer et al. 2015, 5SWS and 5SWZ Gras et al. 2016). The second dimension of the distance PCA, on the other hand, seemed to partially discriminate between class I and class II complexes. To gain some insight in to which structural features were driving this separation, we looked at the distance vectors that were used for each structure (Figure 2b, left). Both for the *α* and the *β* chains, a shift towards the peptide C terminus was observed with decreasing PC2 values. Four representative fingerprints from the edges of the PCA plot are also shown in which the inverted orientation of 4Y19 and 5SWS as well as the shift towards the N terminus for 5TEZ (Yang et al. 2017) are apparent, compared to 3RGV (Yin et al. 2011). In agreement with Figure 1c, class II complexes tend to have higher PC2, which is associated with a shift towards binding at the N terminus of the peptide. 3RGV, which segregates with the class II complexes, is actually a class I complex. Interestingly, however, the YAe62 TCR in the 3RGV complex is reported by the authors to bind both class I and class II complexes with similar orientations, which might explain its positioning with other class II complexes. Strikingly, the other class I complex found with high PC2 is 4JRY, which is also reported to bind with unusual position on top of the N-terminus of the peptide, rather than centrally, where the peptide bulges out (Liu et al. 2013).

**Figure 2:**
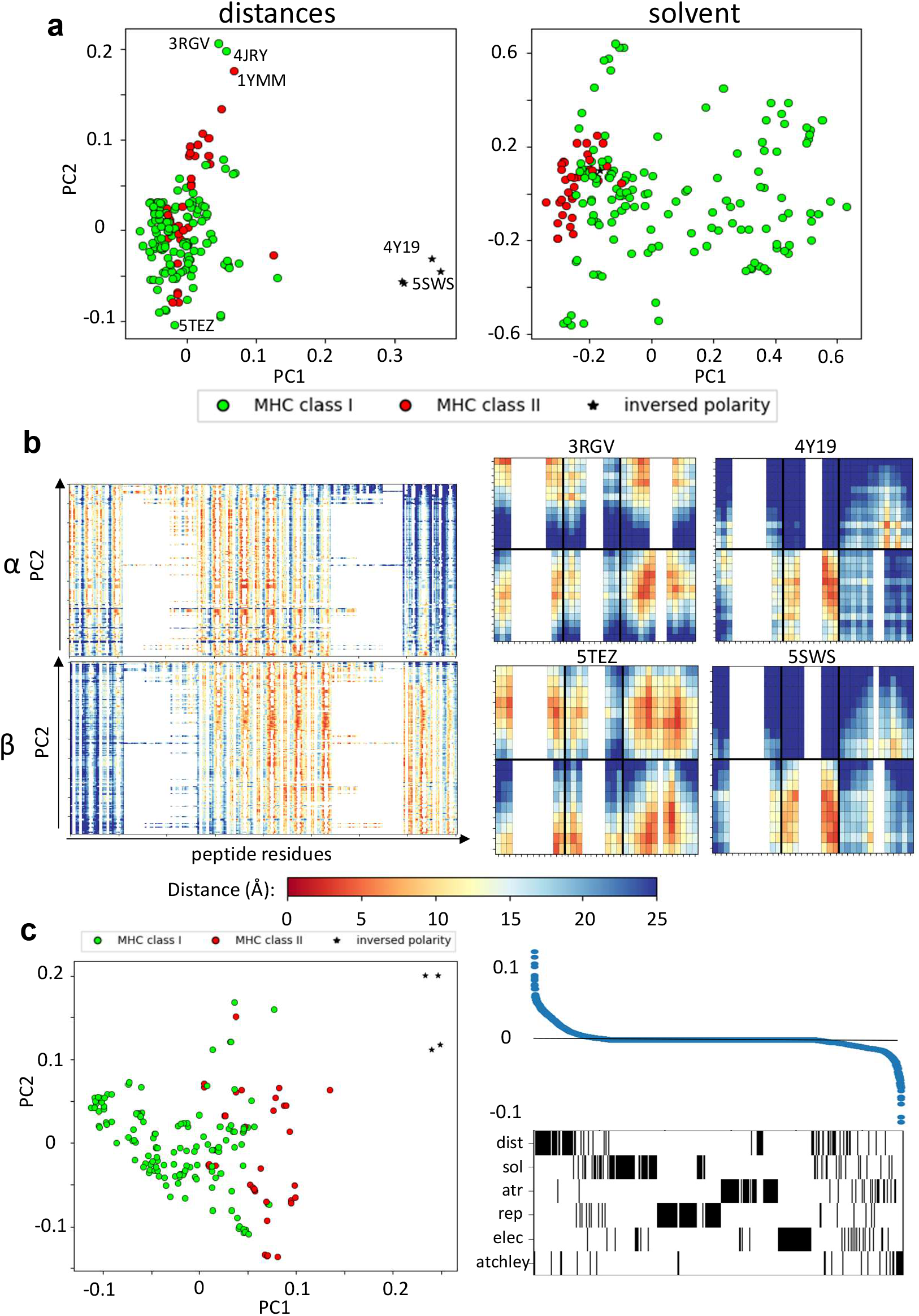
Structural features identify different binding modes. **a.** PCA performed on distances and on solvent energies can separate class I and class II complexes (green and red, respectively). The stars indicate the structures that have been reported to have inversed polarity (i.e. the TCRs bind the pMHC complex at 180 degree angle). Annotated on the distance plot, the structures at the extremes that we analyse in b. **b.** Left: linearised vectors used for the distance PCA, ordered according to their PC2 score. On the x-axis, the minimum distance between each CDR residue and each peptide residue (27-1, 28-1,…,116-1, 117-1, 27-2,…,117-20). Right: fingerprints for 4 representative structures labelled in panel **a** (3RGV high PC2, 5TEZ low PC2, 5SWS and 4Y19 high PC1). **c.** Left: PCA of all feature sets combined, which also shows separation along PC1. Right: loading coefficient of each feature on PC1 and below a barcode to show which set the feature belongs to.

A similar analysis was done on the solvent energy vectors (Figure 2). The PCA suggested a segregation between class I and class II complexes along PC1, although significant overlap was also observed. We therefore looked at what features could be driving the separation along the PC1 (Supplementary Figure S2b). The only evident trend was that all the complexes with high PC1 show a strong unfavourable interaction between the *β* chain and the peptide C terminus (blue in the heatmap). As solvent energy is positive (i.e. unfavourable) when a residue is not solvent-exposed, this suggests that the complexes with higher PC1 make an interaction between the beta chain and the C terminus of the peptide.

Finally, all distance and energy feature sets were combined in a single PCA plotted in Figure 2c (left). Here, the structures with inverted polarity have high PC1, followed by MHC class II complexes and on the left-hand side of the plot are the class I complexes. The loadings of each feature in the set were calculated and the features ranked by loading value (Figure 2c, right). Most of the features which had absolute values greater than 0 (i.e. positive or negative), belong to the distance, the solvent energy or to the Atchley factors datasets, suggesting that these have the strongest discriminatory power.

Overall, these results gave us confidence that meaningful information about the binding interface could be extracted with our pipeline.

### 4.2 Structural information from homology modelled structures cannot distinguish binding pairs in unsupervised settings

We next investigated whether given independently a TCR and a pMHC, we could determine whether we could discriminate between TCR-pMHC interactions in which the TCR binds its cognate antigen and those which do not allow effective binding. The parameters characterising non-binding interactions could obviously not be obtained directly from known structures, since by definition these TCRs would not form stable complexes with the pMHC. We therefore predicted structures for TCR-pMHC combinations by homology modeling using TCRpMHCmodels (Jensen et al. 2019). The pipeline takes a fasta file with a TCR, a peptide and a class I MHC, predicts its three dimensional structure and extracts pairwise distances and binding energies for the interface. The actual sequences are also captured in the form of vectors of Atchley factors as described in the methods.

Because we needed to rely on a structure prediction method, we first evaluated the difference between the features extracted from the original crystallographic structures and from their respective modelled structures (Figure 3 and Supplementary Figure S3a). Taking complex 1AO7 as an example, the fingerprints obtained from the original PDB and from the predicted structures were plotted (Figure 3a). The two complexes have RMSD of about 2Å and it can be seen that the contacts seem to be slightly shifted towards the N terminus of the peptide in the predicted structure compared to the crystal. However, the two fingerprints did not look drastically different.

**Figure 3:**
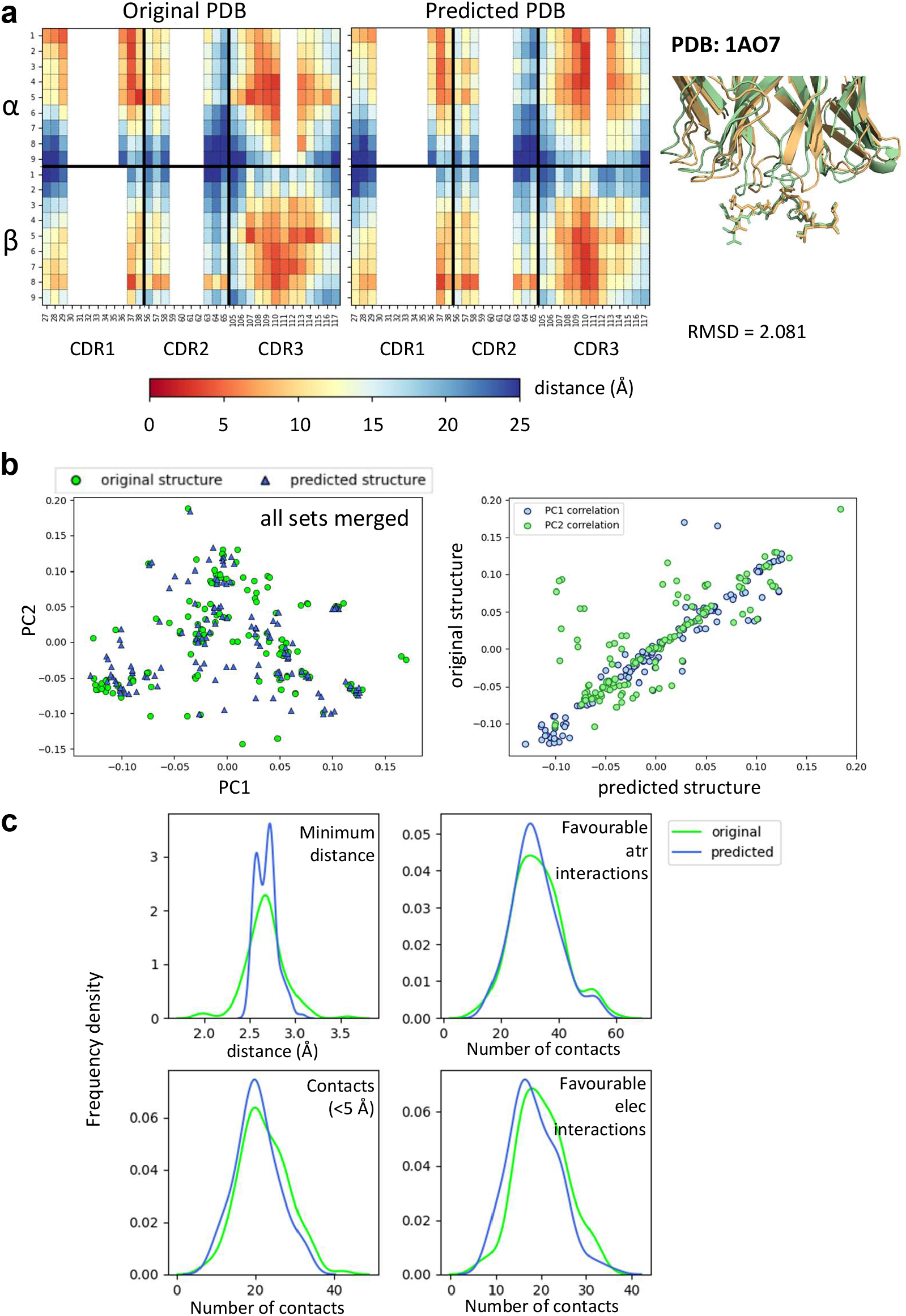
Comparisons between crystal structures and homology predicted structures. **a.** Comparison of fingerprint between the original 1AO7 structure and the one predicted by TCRpMHCmodels. On the right, figure showing how the two structures superimpose in cartoon form (green = original, gold = predicted). MHC not shown for clarity. **b.** Left: PCA on all feature sets showing the difference between crystal structures (green circles) and predicted structures (blue triangles). Right: correlation for PC1 and PC2 values between original and predicted structures. Each blue dot is a complex and has (x,y) coordinates that depend on PC1 values for predicted and original structure. Similarly for PC2 (green dots). PCA for other feature sets in Supplementary Figure S3a. **c.** Frequency distributions of 4 characteristics of the TCR-pMHC complexes comparing the distribution between original and predicted structures. Minimum distance: minimum distance between TCR and peptide; Contacts: number of TCR-peptide residue pairs that are less than 5A apart; Favourable atr/elec interactions: number of favourable (energy *<* 0) interactions between TCR and peptide.

When combining all feature sets and looking at all structures available by PCA, no systematic difference was found between modelled and original structures (Figure 3b and c and Supplementary Figure S3a). There was reasonably good matching between the crystal strucutres and their homology models, although TCRpMHCmodels failed to predict non-canonical binding models. We also compared the distributions of some of the structural features (minimum distance between peptide and TCR, number of contacts and number of favourable interactions), and in general found reasonably good agreement between models and structures. As homology modelling gave us reliable predictions and was necessary to create our negative examples, we decided to use modelled structures for both binding and non-binding complexes, in order to avoid introducing systematic bias.

To create a set of non-binders, a set of shuffled TCR-pMHC complexes from the STCRDab was used (Figure 4a). We then asked whether the structures predicted for non-binders could be discriminated from the binders.

**Figure 4:**
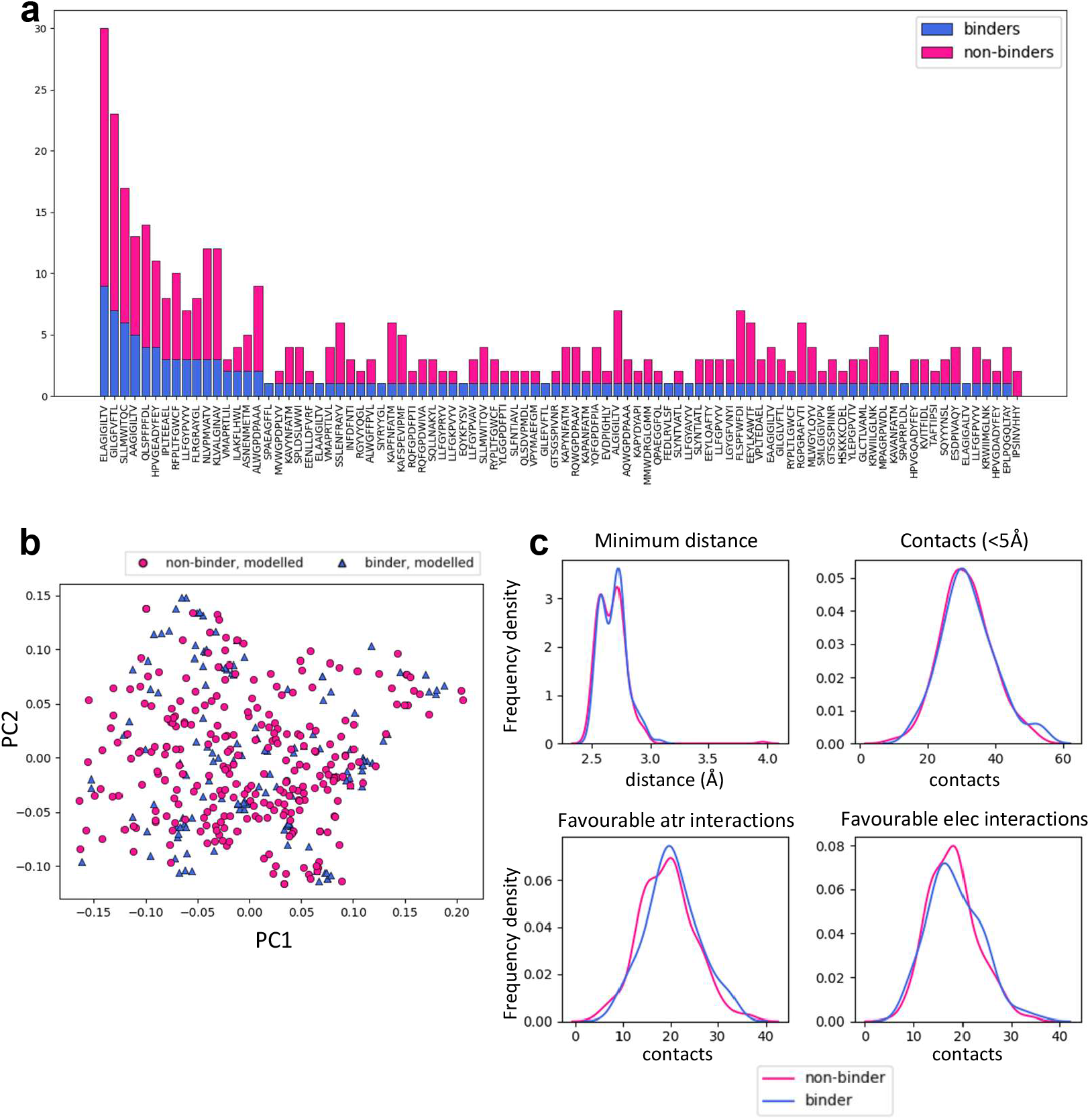
Homology modelled binding and non-binding TCR-pMHC complexes can not be discriminated by PCA. **a.** Summary of the number of STCRDab derived binding and non-binding structures which were modelled. For each peptide in the set, the barplot shows the number of models of binding and non-binding TCRs (blue and magenta, respectively). **b.** PCA of all sets combined showing no separation between binding and non-binding TCR/pMHC homology models. The PCAs for each feature set separately are in Supplementary Figure S3b. **c.** Frequency distributions of 4 characteristics of the TCR-pMHC complexes comparing the distribution between binding and non-binding models. Minimum distance: minimum distance between TCR and peptide; Contacts: number of TCR-peptide residue pairs that are less than 5A apart; Favourable atr/elec interactions: number of favourable (energy *<* 0) interactions between TCR and peptide.

Strikingly, there was no dsicernible separation of binders and non-binders on unsupervised PCAs with any of the distance or energy sets of features (Figure 4b and Supplementary Figure S3b). Basic metrics such as the minimum distance between TCR and peptide and the number of contacts showed similar distributions for binders and non-binders (Figure 4b).

### 4.3 Structural information can discriminate between binders and non-binders using supervised learning

We turned to supervised machine learning methods to try and better discriminate between binding and non-binding pairs. We explored multiple kernel learning (MKL) to combine information from the different feature sets extracted from the modelled interaction surfaces using the pipeline explained above. To assess the potential of our method, a model was trained and tested by cross-validation, using predicted structures derived from the STCRDab, creating a dataset of positives and negatives as described in the methods. Figure 5a and c show the results of 10-fold cross-validation when each different feature set is used separately. Whilst Atchley factors provide the single strongest predictive power (average ROC AUC of 0.763), similar discrimination can be obtained by using distances only (ROC AUC of 0.755), followed closely by attractive van der Waals forces (atr, ROC AUC of 0.74) and solvent energies (ROC AUC of 0.701). The other energetic terms generally showed poorer performance and were excluded from further analysis.

**Figure 5:**
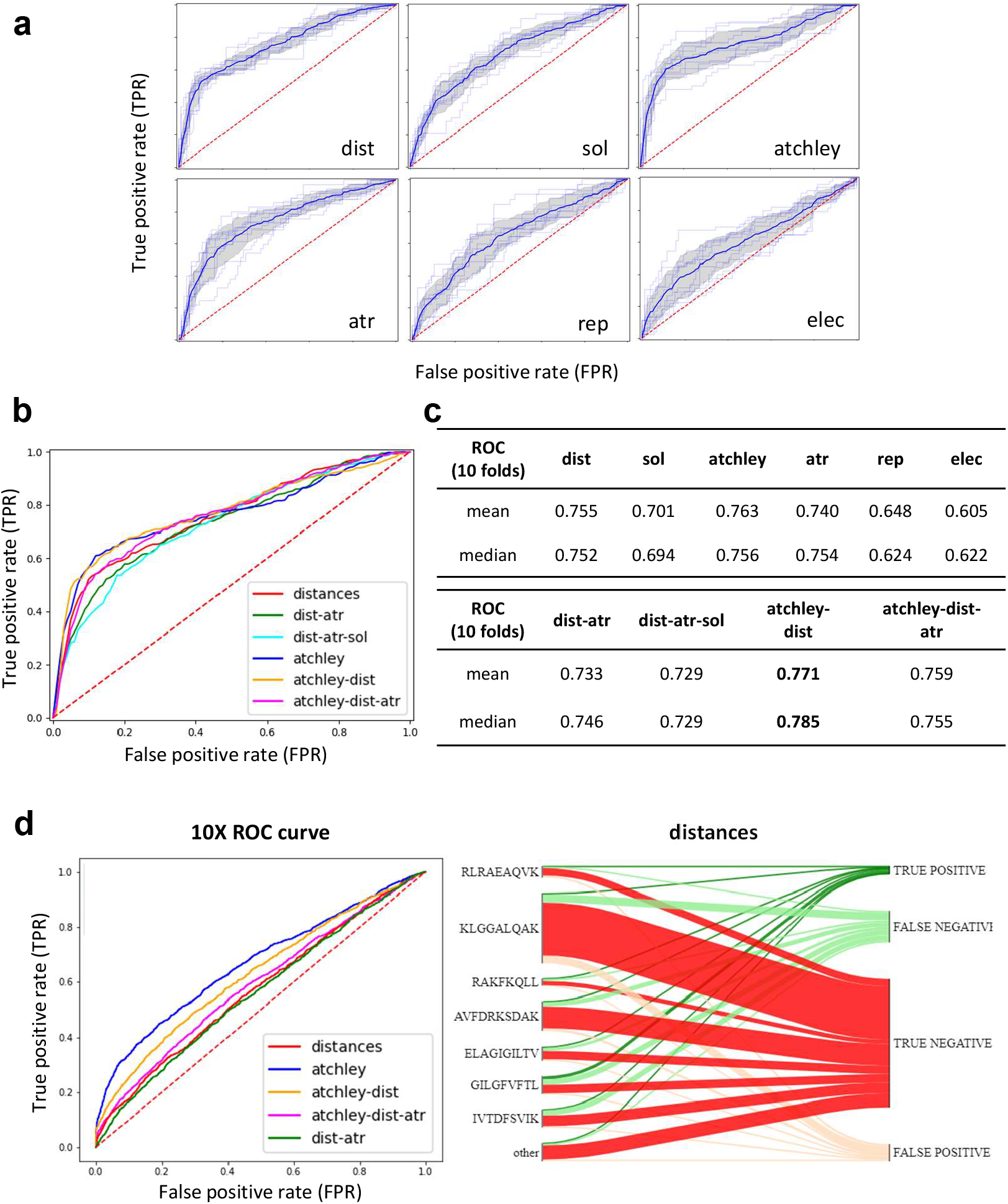
A discriminative classification model can be trained using extracted structural features. **a.** ROC AUC curves of 10-fold CV on the STCRDab training set with each feature set separately. The faint line are the results for each individual fold, whilst the dark line represents the interpolated average results, with the shaded area as the standard deviation. **b.** Interpolated ROC AUC curves for 10-fold CV obtained when combining different feature sets for prediction. **c.** Tabular results for curves showed in a. and b.. **d.** Left: ROC curves obtained when the model trained on the STCRDab set is used for prediction on the 10XGenomics validation set. Right: for the model trained on STCRDab using the distance dataset only, the diagram shows which proportion of examples from each epitope are classified correctly (true positives and true negatives) or incorrectly (false positives and false negatives).

We next combined the feature sets to create a single classifier (Figure 5b and c). Using Atchley factors, distances and attractive van der Waals forces together achieved a similar performance to using each set of features independently, whilst combination of Atchley factors and distances only gave a slight increase in performance compared to each of the two sets separately. Interestingly, although performance did not change much in this more complex model, the weights assigned to the kernels constructed for each feature set were similar, suggesting that no single feature set was more important than the others in the overall model.

We then went on to validate the trained model on the other 5 datasets described in the methods. Because we wanted to test how generalisable the rules that the classifier had learnt were, we did not train the classifier again on the new sets, but used the model trained on the STCRDab set to predict the new complexes. Results from validation are presented in Figure 5d and Supplementary Figure S4 and summarised in Table 1. Overall, the models with the highest ROC AUC consistently included sequence information. Moreover, addition of structural features often did not improve predictive power. However, structural features often allowed some level of discrimination, independently of the sequence information, suggesting that the model might be learning something about the binding modes of these complexes. Interestingly, the models which used structural features had consistently higher recall.

**Table 1:** Results of out-of-sample validation. Results of predicting the validation sets with the model trained on the STCRDab set, using different subsets of features. In each section, the best-performing model is highlighted in bold and underlined.

| set | % pos | combo | roc | avg<br>precision | accuracy | precision | recall |
| --- | --- | --- | --- | --- | --- | --- | --- |
| **B10x** | 21.17 | distances | 0.574 | 0.289 | 0.739 | 0.315 | 0.198 |
|  |  | dist-atr | 0.562 | 0.260 | 0.726 | 0.294 | **<u>0.210</u>** |
|  |  | atchley | **<u>0.668</u>** | **<u>0.441</u>** | **<u>0.805</u>** | **<u>0.751</u>** | 0.117 |
|  |  | atchley-dist | 0.629 | 0.375 | 0.786 | 0.487 | 0.166 |
|  |  | atchley-dist-atr | 0.590 | 0.317 | 0.766 | 0.382 | 0.173 |
| **Dash** | 7.34 | distances | 0.591 | 0.114 | 0.757 | 0.116 | **<u>0.350</u>** |
|  |  | dist-atr | 0.645 | 0.123 | 0.802 | 0.139 | 0.326 |
|  |  | atchley | **<u>0.700</u>** | **<u>0.188</u>** | **<u>0.905</u>** | **<u>0.209</u>** | 0.107 |
|  |  | atchley-dist | 0.599 | 0.175 | 0.798 | 0.133 | 0.318 |
|  |  | atchley-dist-atr | 0.645 | 0.146 | 0.824 | 0.153 | 0.309 |
| **expt** | 12.70 | distances | 0.727 | 0.326 | 0.714 | 0.262 | **<u>0.688</u>** |
|  |  | dist-atr | 0.709 | 0.423 | 0.667 | 0.205 | 0.563 |
|  |  | atchley | 0.816 | **<u>0.704</u>** | **<u>0.825</u>** | **<u>0.393</u>** | **<u>0.688</u>** |
|  |  | atchley-dist | **<u>0.823</u>** | 0.659 | 0.754 | 0.297 | **<u>0.688</u>** |
|  |  | atchley-dist-atr | 0.770 | 0.515 | 0.698 | 0.238 | 0.625 |
| **atlas** | 89.06 | distances | 0.487 | 0.897 | 0.827 | 0.892 | 0.917 |
|  |  | dist-atr | 0.518 | 0.907 | 0.794 | **<u>0.901</u>** | 0.863 |
|  |  | atchley | **<u>0.632</u>** | **<u>0.938</u>** | **<u>0.891</u>** | 0.891 | **<u>1.000</u>** |
|  |  | atchley-dist | 0.551 | 0.918 | **<u>0.891</u>** | 0.891 | **<u>1.000</u>** |
|  |  | atchley-dist-atr | 0.547 | 0.916 | 0.865 | 0.896 | 0.960 |
| **newVdj** | 0.72 | distances | 0.521 | **<u>0.010</u>** | 0.865 | 0.010 | **<u>0.186</u>** |
|  |  | dist-atr | 0.521 | 0.008 | 0.896 | **<u>0.013</u>** | **<u>0.186</u>** |
|  |  | atchley | **<u>0.570</u>** | **<u>0.010</u>** | **<u>0.987</u>** | 0.000 | 0.000 |
|  |  | atchley-dist | 0.541 | **<u>0.010</u>** | 0.954 | 0.009 | 0.047 |
|  |  | atchley-dist-atr | 0.546 | 0.008 | 0.947 | 0.000 | 0.000 |

The ATLAS proved to be a very hard set to predict overall. This might be due to each complex being only a few mutations away from the crystal structure deposited in the PDB, which might have on one hand made the modelling easier, but on the other hand made it harder for the classifier to tell the difference between a binding and a non-binding pair which differ at only one amino acid. Moreover, some of the included mutations occur at the MHC, which is not considered when extracting features. Finally, the ATLAS set does not have a strict definition of binding, as for the other sets which derive from tetramer-sorting experiments, but rather the complexes show a range of affinities, and it is hard to define a strict threshold to define binding.

### 4.4 Classifier performance varies between epitopes

A known hard task for a classifier trained on a small subset of the epitopes that our immune system is exposed to, is to generalise to epitopes not present in the training set. It is apparent from the diagrams showing mis-classification in Figure 5d (right) and Supplementary Figure S4b that some peptides were indeed easier to classify than others. Figure 6a shows the classifier performance on 4 representative epitopes. For a perfect classifier, the decision score for positive and negative samples (equivalent to the distance of a point from the decision hyperplane in the case of an SVM) should have non-overlapping distributions. However, for peptide antigen AVFDRKSDAK the distributions for binding and non-binding TCRs almost completely overlap, suggesting that the classifier has not learnt useful information from the data. For peptide LLFGYPVYV, on the other hand, the separation between the two groups of TCRs is almost perfect. The classification of TCRs specific for the ELAGIGILTV and ASNENMETM peptides showed an intermediate pattern. Overall, the classification of TCRs for different epitopes show very significant differences in performance, (Figure 6b), as has been observed previously for other models (Moris et al. 2020). This also suggests that the overall performance as showed in Table 1 is somewhat misleading, as it will be skewed by the more abundant epitopes.

**Figure 6:**
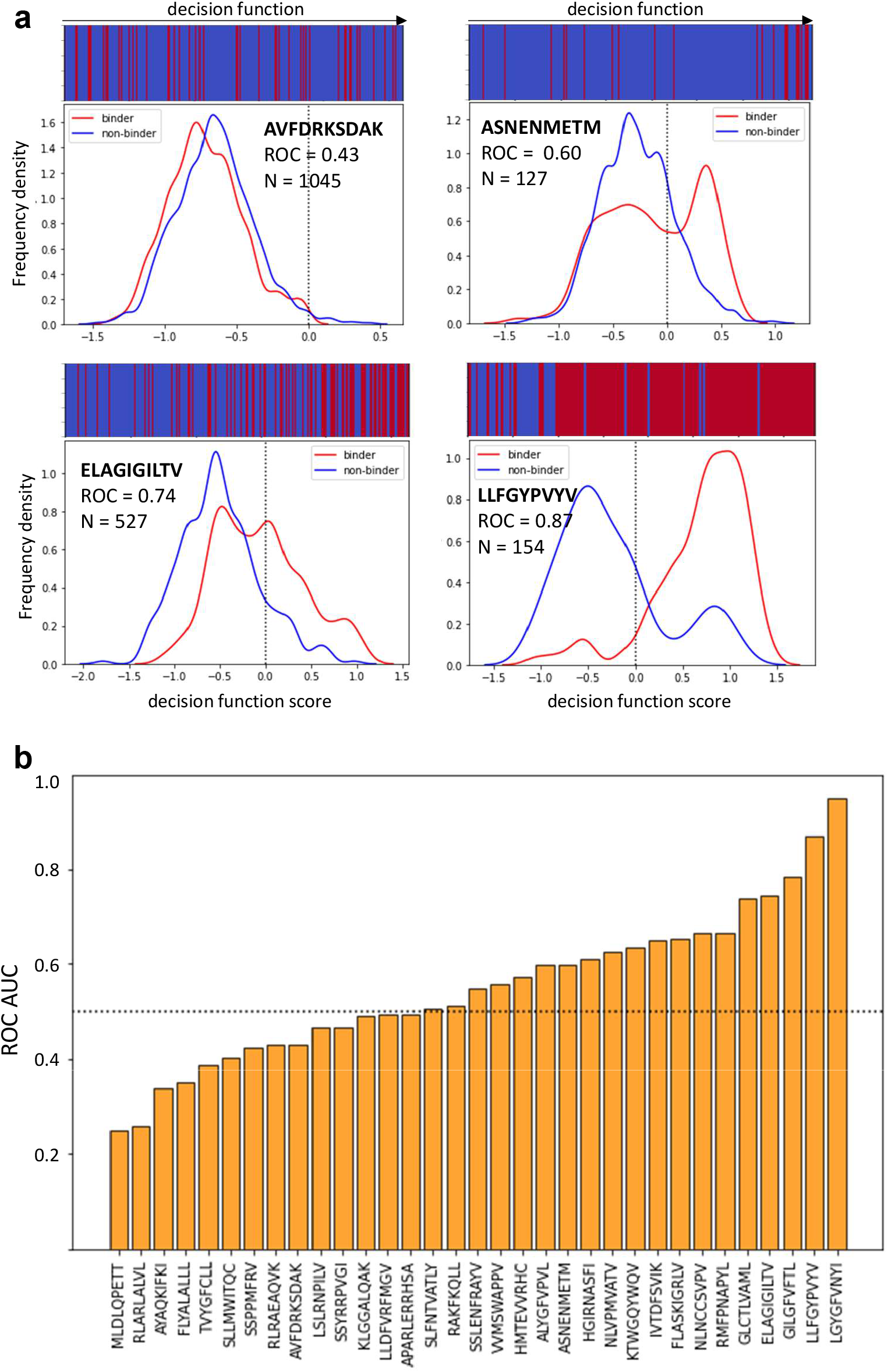
The performance of the model is pMHC dependent. **a.** Examples of 4 different epitopes. The frequency distributions of model decision function scores (for an SVM, this correponds to the distance from the separating hyperplane, drawn as a dotted line) for binding and non-binding TCRs recognising each epitope. The bar at the top shows the order in which binding and non-binding examples appear when ranked by decision function. For good classification, the bar should be mostly blue on the left and mostly red on the right. **b.** The bar plot shows ROC AUC for all peptides which have at least 2 positive and 2 negative examples. This data comes from concatenating the predictions for all the validation sets when Atchley factors, distances and attractive van der Waals forces are used.

### 4.5 Homology modelling performance impacts classifier performance

We wondered whether the difference in performance could be due to the performance of the homology modelling tool used. For each structure, we retrieved the information about the sequence similarity between the structure of interest and the template used to model it. We then plotted the classifier performance as a function of sequence similarity (Figure 7a).

**Figure 7:**
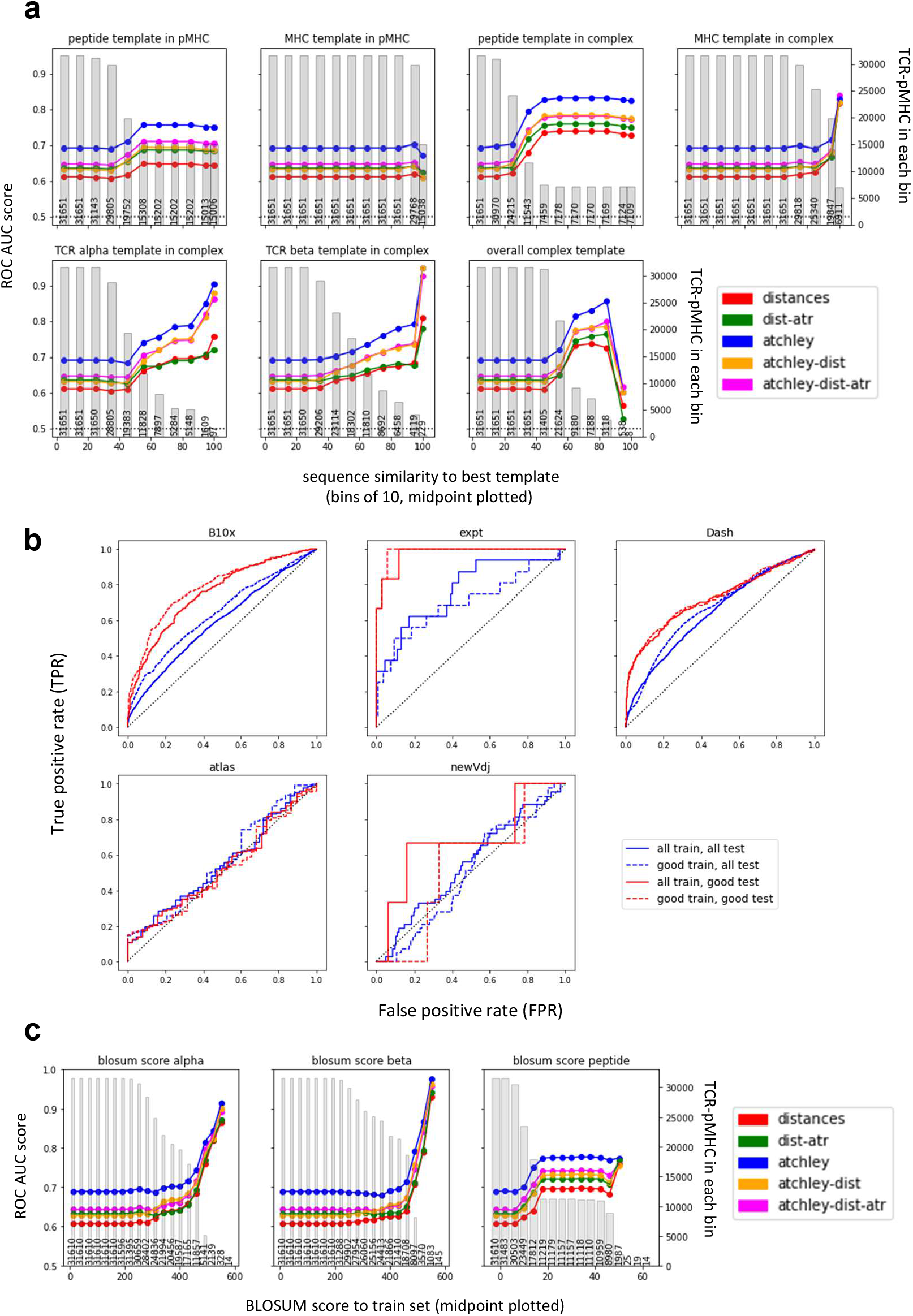
Classifier performance is dependent on sequeunce homology of the target TCR-pMHC. **a.** The performance from all validation sets were combined, and stratified by the similarity between the sequence of the target complex to be classified and the relevant homology modelling template (as outputted by TCRpMHCmodes and outlined in Jensen et al. 2019). Mean performance (ROC AUC) in each range of homology is calculated and plotted at the range midpoint. The grey bars show the number of structures that contribute to the performance for each point. **b.** Performance of each of the validation set when the model is trained on the entire STCRDab set (all train) or only the STCRDab structures with good templates (as defined in methods - good train), and when predictions are made on all complexes (all test) or only complexes with good templates (good test). **c.** Equivalent analysis to a. but calculating the BLOSUM score between each example and the closest example in the train set, for each chain separately. The higher the BLOSUM score, the more similar the sequence is to one found in the training set. In each plot, the grey bars show the number of structures in each bin.

Overall, there was a trend for better templates (increased sequence similarity) to correlate with better classifier performance (observed as an increase in performance to the right of the individual panels). Interestingly, however, the same trends were observed also when classification was based only on sequence information suggesting that this might not be related only to the accuracy of the homology modelling. The templates for the homology modelling and the training set for our classifier are overlapping sets (as both are using the complexes for which a crystal structure is available) and our results might be reflecting the increased density in the feature space of known complexes. To investigate this, we also computed the BLOSUM scores from the train set for all the complexes we predicted (Figure 7c). Indeed, a decrease in classifier performance is observed when the BLOSUM score decreases, i.e. when the TCR-pMHC pair that we are trying to predict is less similar to the training set pairs. Interestingly, in all cases the performance of the classifier is more dependent on TCR homology, than on peptide homology. It is important to note that the observed relationship between classifier performance and sequence homology allow us to predict *a priori* which TCR/peptide binding predictions will carry greater confidence. In fact, by considering the epitope and complex homology templates, we are able to select *a priori* a subset of structures on which our model will perform better (Figure 7b).

### 4.6 Effect of affinity on the predictor

Because the classifier relies on structural information and it is trained on the set of TCR-pMHC pairs that have a known crystal structure, we wondered whether the model could predict binding affinity as well as a binary binding/non-binding classification or whether higher decision function scores were assigned to higher-affinity complexes (i.e. whether complexes which bind with high affinity are called binders with higher confidence). To address this, the TCR-pMHC pairs from the ATLAS (Borrman et al. 2017) were retrieved and their score predicted. The score for each complex was then correlated (Spearman) to their measured affinity, removing all complexes with undetectable binding and adjusting the ΔG and *K_D_* as in the original publication (Table 2). Unexpectedly, the only significant correlation was between sequence features (Atchley factors) and *k_off_*. The model therefore does not successfully capture the structural information which determines the affinity of the complex and its performance is not biased towards detection of high-affinity pairs.

**Table 2:** Correlations of affinity metrics and decision function scores. Spearman correlation is calculated for each affinity metric for predictions made for each of the models trained.

|  | distances |  | dist-attr |  | atchley |  | atchley-dist |  | atchley-dist-attr |  |
| --- | --- | --- | --- | --- | --- | --- | --- | --- | --- | --- |
|  | Spearman | p-value | Spearman | p-value | Spearman | p-value | Spearman | p-value | Spearman | p-value |
|  | R |  | R |  | R |  | R |  | R |  |
| $K_D$ ($\mu\text{M}$) | -0.076 | 0.188 | -0.057 | 0.322 | -0.006 | 0.914 | -0.048 | 0.402 | 0.154 | 0.099 |
| $k_{on}$ ($\text{M s}^{-1}$) | 0.126 | 0.177 | 0.153 | 0.101 | 0.084 | 0.371 | 0.173 | 0.063 | 0.050 | 0.592 |
| $k_{off}$ ($\text{s}^{-1}$) | 0.056 | 0.551 | -0.077 | 0.412 | **0.277** | **0.003** | 0.106 | 0.260 | -0.070 | 0.221 |
| $\Delta G$ (kcal/mol) | -0.080 | 0.167 | -0.065 | 0.258 | -0.022 | 0.702 | -0.060 | 0.338 | -0.070 | 0.221 |

### 4.7 Benchmarking against existing tools

Finally, we compared the performance of our classifier against the recently published ERGO (Springer et al. 2020) and ImRex (Moris et al. 2020, Table S1). Both ERGO and ImRex were trained on the VDJdb set (Bagaev et al. 2020), as described in the original publication, rather than the much smaller set of binder used by our algorithm. The trained models are available as an online tool for ERGO (http://tcr.cs.biu.ac.il/) and on GitHub for ImRex (https://github.com/pmoris/ImRex).

The classifiers were all tested on the same set of binder and non-binder TCR-pMHC sets. Figure 8 and Supplementary Table S1 show the results divided by peptide. The results are organised in 3 scenarios depending on whether the peptide is present in neither, either or both of the train sets.

**Figure 8:**
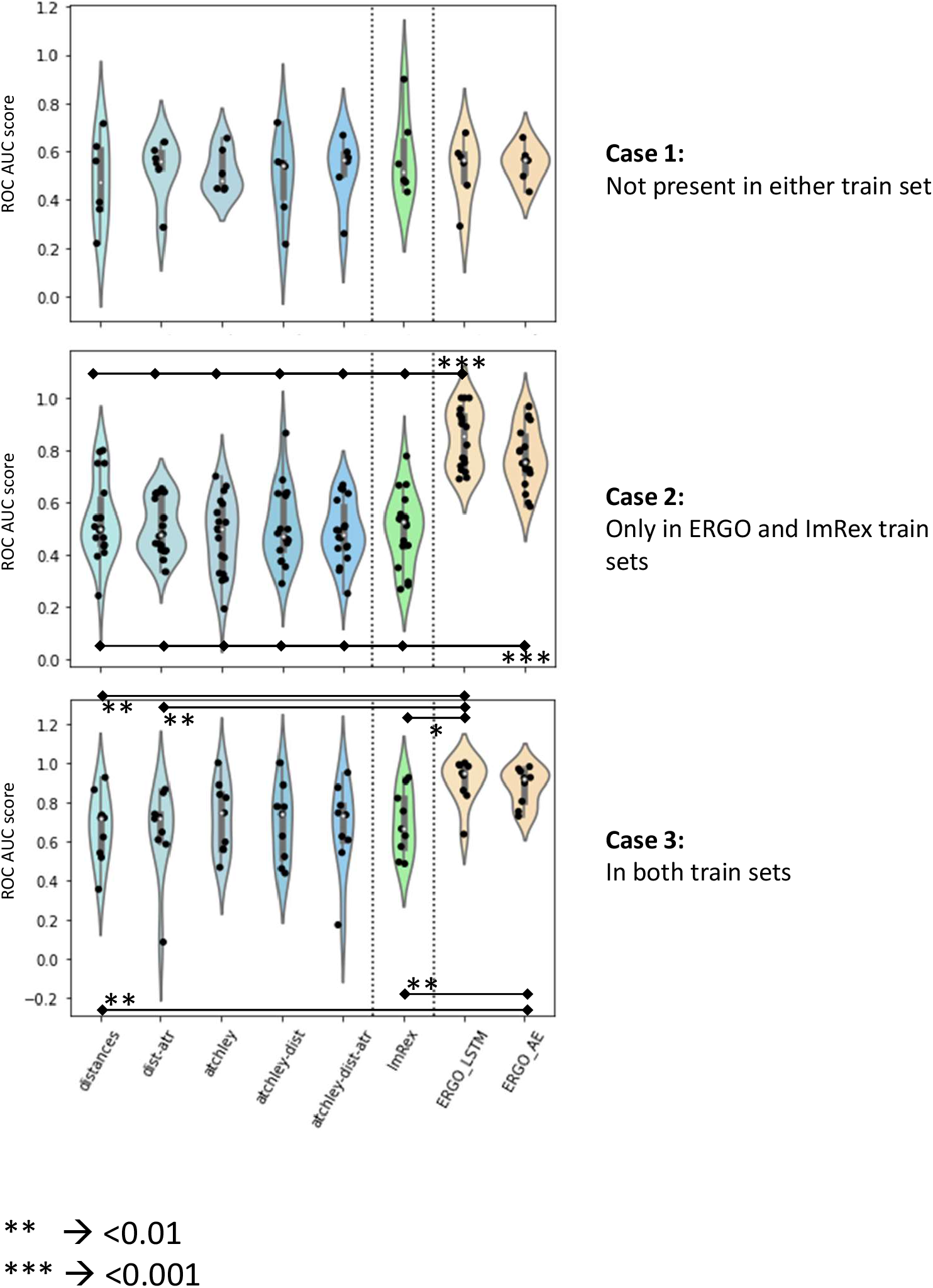
Comparison of performance with other published tools. In each violin plot, a dot is an epitope for which performance is calculated. In Case 1, only epitopes that are not present in the PDB or in the VDJDb train sets are included. In Case 2, only epitopes that are present in the VDJDb but not in the PDB are included. In Case 3, only epitopes which are in both training sets are included. Significance values are shown by asterisks.

When compared on epitopes that are not present in either train set (Case 1), all the models perform in a similar manner. Interestingly, none of the sequence-based classifiers outperforms the structure-based classifier. When the epitopes are present in the VDJDb but not in the STCRDab (PDB) set (Case 2), both ERGO models significantly outperform all other models in prediction, including ImRex. Finally, when peptides are present in both train sets (Case 3), ERGO outperforms all models except the ones which include Atchley factors information.

Taken together, these results suggest that the structure-based models developed in this study perform as well as the state-of-the-art sequence-based models in predicting binding to novel pMHC, despite learning from a much smaller training set.

## 5 Discussion

Previous study of the binding geometry of TCRs to the pMHC complex has been largely focused on measuring the diagonal angle and the orientation of the TCR with respect to the MHC. In the present study, a number of different features were extracted to try and recapitulate both the conformation and the energetic profile of the binding interface. A survey of the crystal structures available showed that, in agreement with Glanville et al. 2017; Ostmeyer et al. 2019, stretches of amino acids at the centre of the CDR3 in the TCR*α* and *β* chains are within contact distance of the peptide. This information was also recapitulated by the energy profiles, suggesting that not only can they interact, but that they make favourable interactions. Although no conserved binding hotspots were detected within the CDR, we were able to identify different binding modes simply from the features extracted.

Conserved binding geometry has been reported in TCRs that bind the same MHC complex (Blevins et al. 2016) and recently Singh et al. 2020 showed that a difference can be detected between pMHC class I and class II binding. Such a difference is also reported in this analysis, and detected both at the conformational level (in terms of pairwise distances) and at the energetic level. As reported by Singh et al. 2020, our analysis also showed that TCRs binding MHC class I tend to be closer to the C-terminus of the peptide, whilst TCRs binding class II complexes sit more centrally or towards the N-terminus. Moreover, the energetic features suggest that a difference between class I and class II complexes can also be found in the energetic profiles that drive these interactions. As well as the difference between class I and class II, the spatial features extracted from the structures were readily able to distinguish TCRs which bind with reversed polarity to the pMHC complex, as described by Gras et al. 2016 and Beringer et al. 2015, and identify class I complexes with different non-canonical binding modes to the peptide (Yin et al. 2011; Liu et al. 2013). This suggests that the features extracted are informative of the biology of this system.

The information collected from these structures was also sufficient to build a classifier able to discriminate between TCR-pMHC binding from non-binding pairs. The generalisability of the classifier was tested on multiple independent datasets, collected and analysed independently. Physical interaction features on their own proved sufficient to distinguish binding and non-binding complexes to a similar degree to published tools which are based on sequence information alone (Figure 8). Interestingly, merging of sequence and physical features in the same model did not improve the performance in terms of ROC AUC, although often improved the recall of the sequence-based model. This is an important characteristic, as in real-life applications a classifier like the one presented could be used to screen candidate TCRs against an epitope of interest, for example with the aim of identifying tumour-infiltrating lymphocytes that can recognise tumour neoantigens. In this context, *in-silico* screening would be followed by experimental validation. Because the events of interest are a very small number compared to the total number of events (i.e. binders *<<* non-binders), it would be more important to correctly classify more of the binders than of the non-binders, i.e. a higher number of false positives, which can be screened out during experimental validation, would be less problematic than a higher number of false negative, which would not be experimentally validated.

Compared to other published classifiers (Glanville et al. 2017; Dash et al. 2017; Tong et al. 2020), the classifier presented here is different in that it does not need to be trained on a known subset of TCRs recognising a specific peptide to be able to predict more binders, but rather it can learn from any set of TCR-pMHC pairs already available and generalise what it has learnt to the problem at hand. This suggests that there are conserved features to the TCR-pMHC interface which can be learnt and used for prediction. ERGO and ImRex (Springer et al. 2020; Moris et al. 2020) have pioneered this approach, although they only focussed on information that can be extracted from the sequence. ImRex is a bit more similar to the classifier presented, as it encodes the binding interface using amino acid characteristics rather than pure sequence encoding. Of note, all of the results that we have presented here use the model originally trained on the STCRDab set, which was never re-trained on the new sets of structures. This is not the case for other published tools, which achieve better discrimination but only after training on a section of the validation set.

We extended the approach adopted by ImRex and decided to rely on the structure of teh whole TCR-pMHC complex. Modelling of mutations within the existing crystal structures has recently proved a successful approach to ranking candidate peptide epitopes from a phage screen against target TCRs (Borrman et al. 2020). Here, we see from the weights assigned to each combined kernel that the physical interactions encoded by the distances and the attractive van der Waals forces were equally as important as the sequence information, suggesting that physical interactions can be used to predict binding. Moreover, the classifier here presented is trained on about 400 binding and non-binding pairs, which recognise 93 different epitopes. This is a much smaller set than the VDJdb used by ERGO and ImRex (40,000 TCRs and 200 peptides in ERGO and 14,000 CDR3*β* and 118 peptides in ImRex), but achieves similar performances. This might indicate that the information learnt from the structural information is more readily generalised to an unseen case.

As more structures for more diverse epitopes become available, the performance of the classifier may well improve. However, the complex biology of the system will always be a factor limiting performance. For example, if a small proportion of TCRs bound to the pMHC complex with conformations that are significantly different from canonical binding, we might never be able to predict their binding with a tool that has learnt on a subset of canonical TCRs. This may well be the case with other structures with reversed polarity or complexes with unusual binding highlighted in Figure 2a.

Most of the results presented has been based on a binary classification of TCR-pMHC complexes as binding or non-binding. In reality, the interaction between TCR and pMHC is characterised by a graded affinity scale. This is of interest as there are multiple metrics that contribute to overall affinity and are important for T cell activation dynamics - *K_D_*, *k_on_*, *k_off_*, half-life - (Gálvez et al. 2019; Lever et al. 2017; Stone et al. 2009) and it is not yet clear what features in the structure can drive them. No correlation between the classifier score and affinity or kinetic parameters was detected for the ATLAS structures (Borrman et al. 2017). However, the original ATLAS publication showed a correlation between the attractive van der Waal force as calculated by Rosetta (here atr) and the experimentally-measured affinity, similar to the one reported by Erijman et al. 2014 on an unrelated system. Because the affinity is driven by structure, we believe the PDB classifier could also be optimised for rough affinity prediction, although better methods of modelling the mutations into the structures might have to be explored.

Finally, the major difference between this classifier and most of the work published so far is that it relies on an available TCR*αβ* pairs and cannot be used on unpaired chains. This is a limitation to the direct application of the classifier as alpha/beta pairing is typically not available from bulk TCRseq data. However, unpaired *α* and *β* chains only contain a portion of the binding site information, and the assumption that binding of the *β* chain only is sufficient is clearly not true in every case. Carter et al. 2019 show that the information encoded in the *αβ* pair is synergistic, i.e. that the pairing carries more than the sum of the individual chain information. Moreover, their survey of the VDJdb shows instances where the same *α* chain paired with different *β* chains recognise different epitopes, or where CDR3*α* and *β* annotated to bind epitopes from different species come together to bind yet another peptide. Overall, we believe this to be strong motivation to work on *αβ* pairs. Future work will focus on understanding whether candidate *αβ* pairs that bind a specific antigen can be inferred from TCR clones that are expanded during an immune response.

## 7 Supplementary Material

The following are supplied as supplementary materials:

1. Sequences for all the datasets used, specifically:

- **sequences from STCRDab PDB files** - these are the sequences from the PDB files used for the initial feature extraction
- **STCRDab set metadata** - metadata associated with the sequences from the STCRDab
- **10XGenomics set sequences** - sequences for the structures included in the 10X set
- **experimental constructs sequences** - sequences for the structures included in the expt set
- **Dash set** - sequences for the structures included in the Dash set
- **ATLAS sequences** - sequences for the structures included in the TCR ATLAS set, including the affinity information from the ATLAS
- **VDJDb validation sequences** - sequences for the structures included in the new VDJDb set
2. All result files with decision function scores for each TCR-peptide pair. A README file is included with filename explanations.

**Figure S1:**
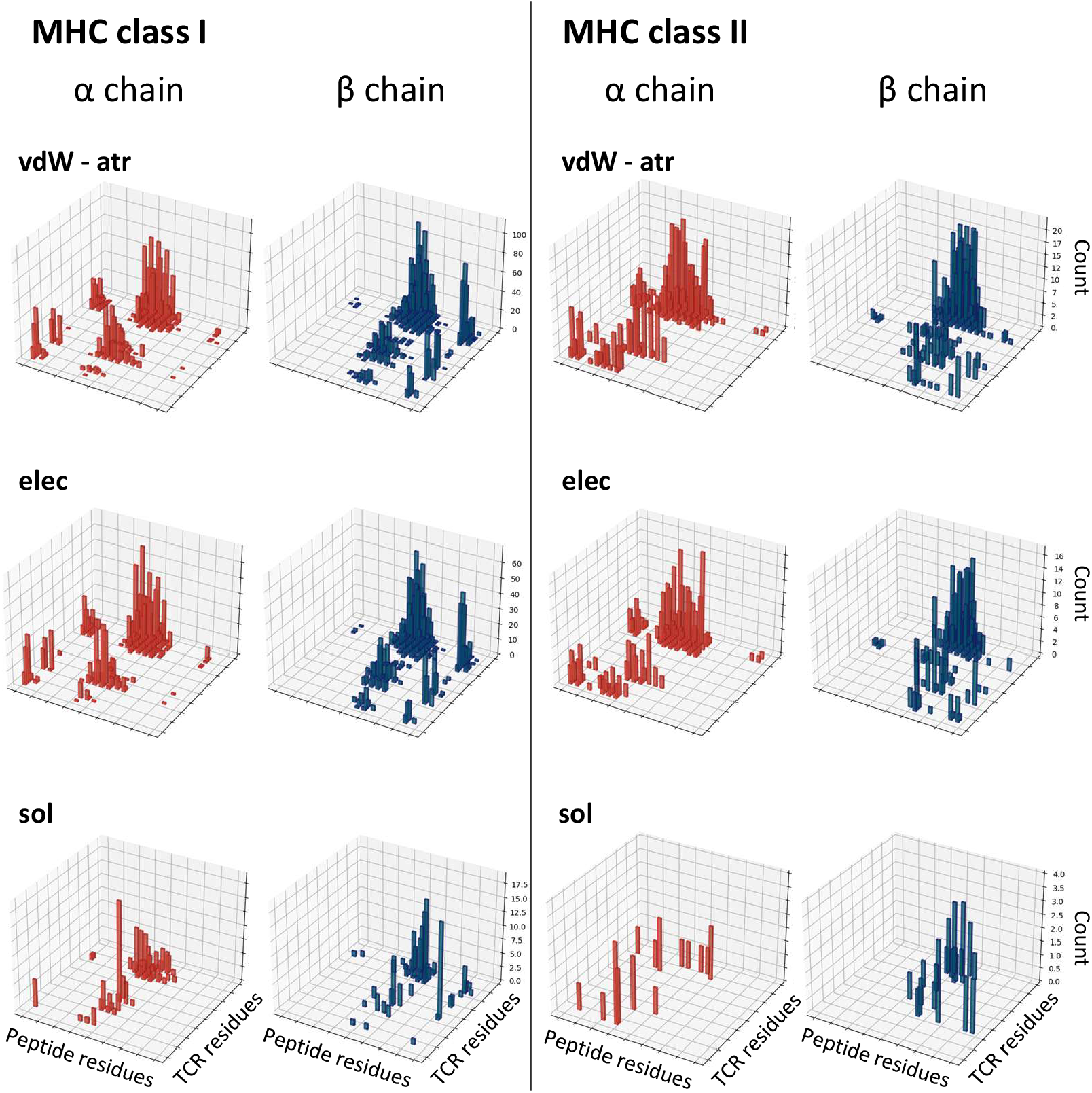
Energy interactions for class I and class II complexes. Analogous to Figure 1c, but for all energy feature sets. The histograms show the number of structures that make a favourable contact (energy *<* 0). Repulsive vdW excluded as this component is always *>* 0.

**Figure S2:**
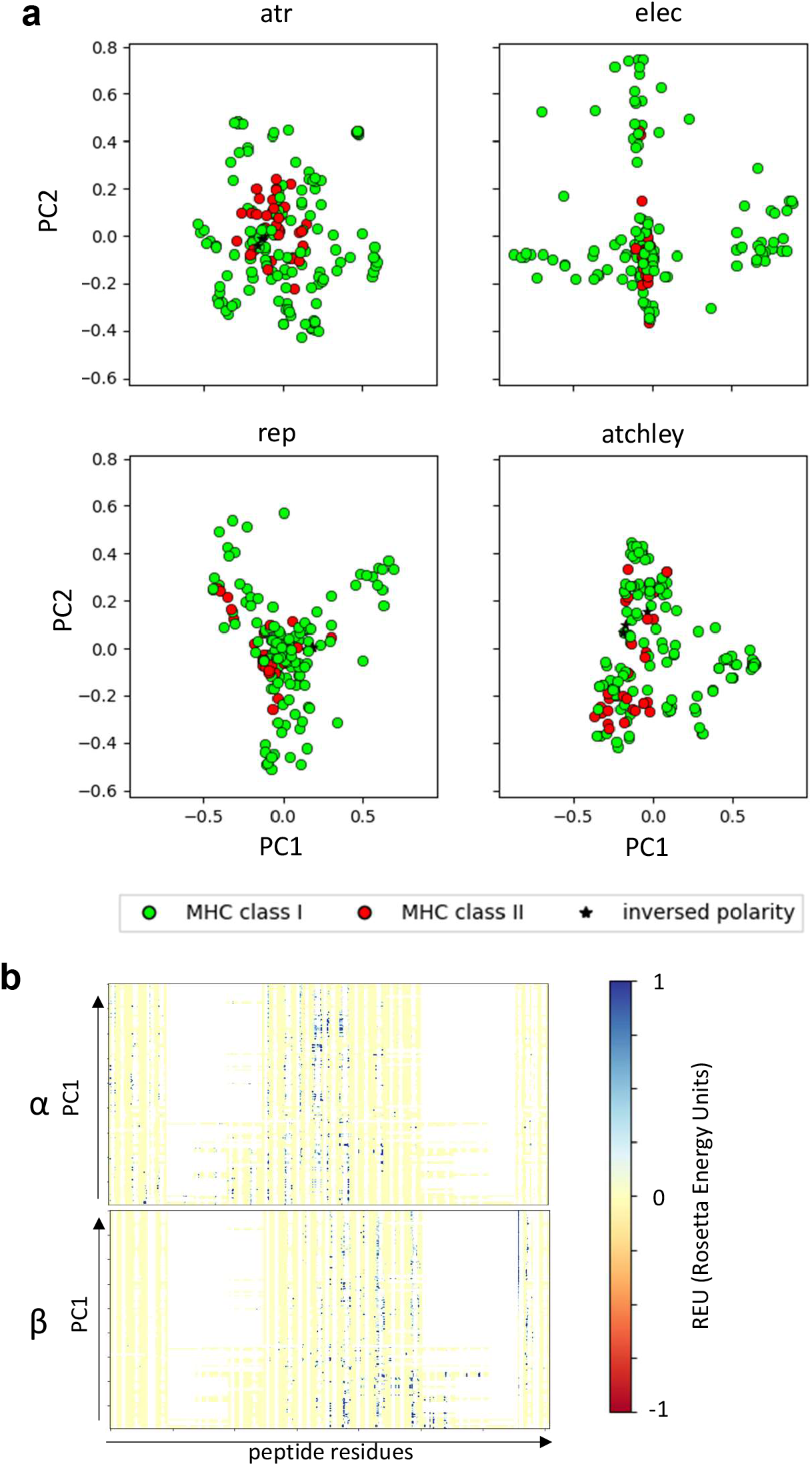
PCA on all extracted features. **a.** PCA for feature sets not included in Figure 2a. Class I and class II complexes are shown in green and red, respectively. The stars indicate the structures that have been reported to have inversed polarity (i.e. the TCRs bind the pMHC complex at 180 degree angle). **b.** Linearised vectors used for the solvent energy PCA, ordered according to their PC1 score. On the x-axis, the calculated solvent energy between each CDR residue and each peptide residue (27-1, 28-1,…,116-1, 117-1, 27-2,…,117-20). Analogous to Figure 2b.

**Figure S3:**
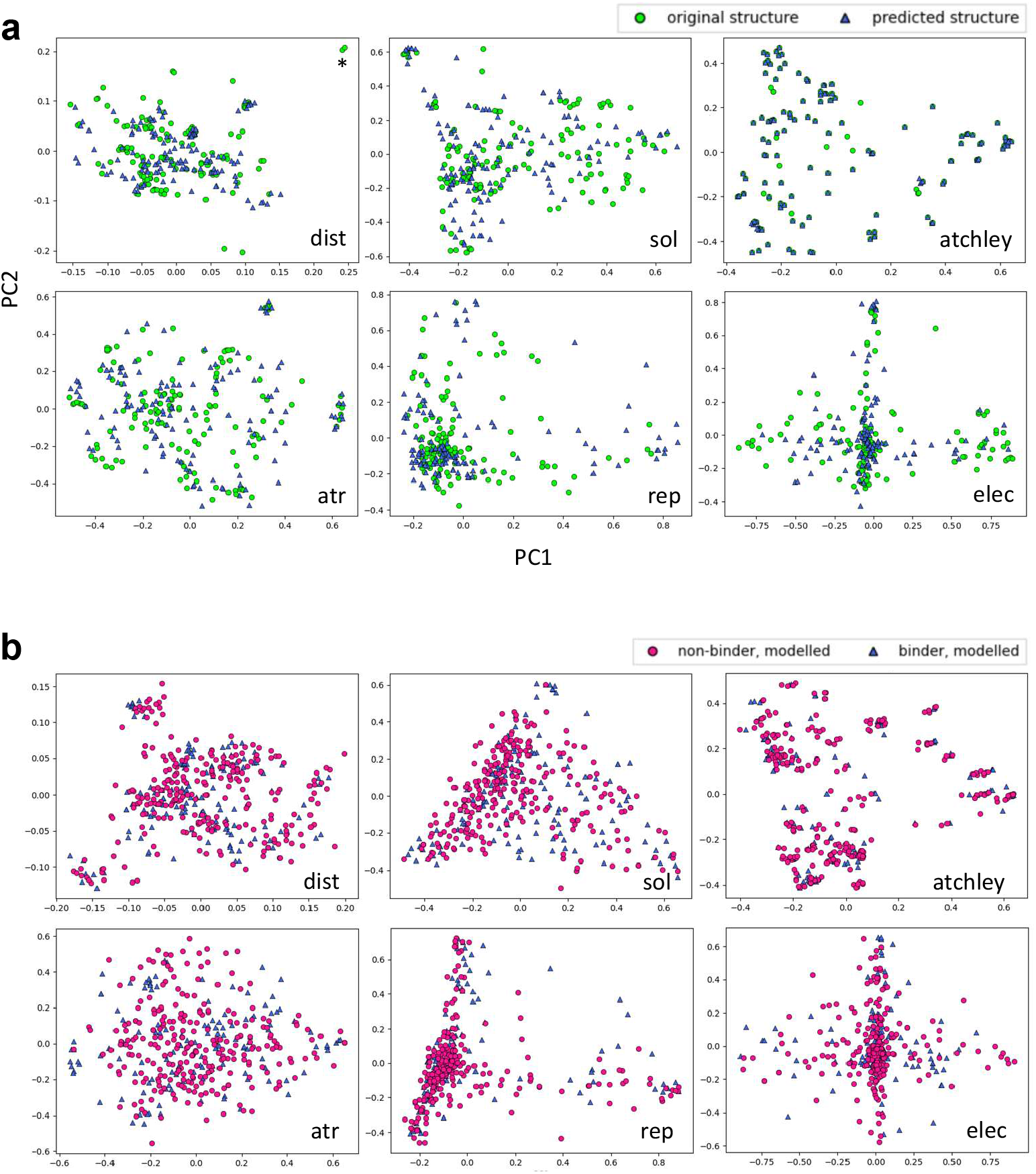
PCA of original vs predicted and of binding vs non-binding. **a.** PCA for each set showing overlay between original and predicted structures. Asterisks (*) in the distance plot indicates the inversed polarity structures. **b.** PCA for each set showing overlay of binding and non-binding complexes (predicted structures, blue triangles and magenta circles, respectively).

**Figure S4:**
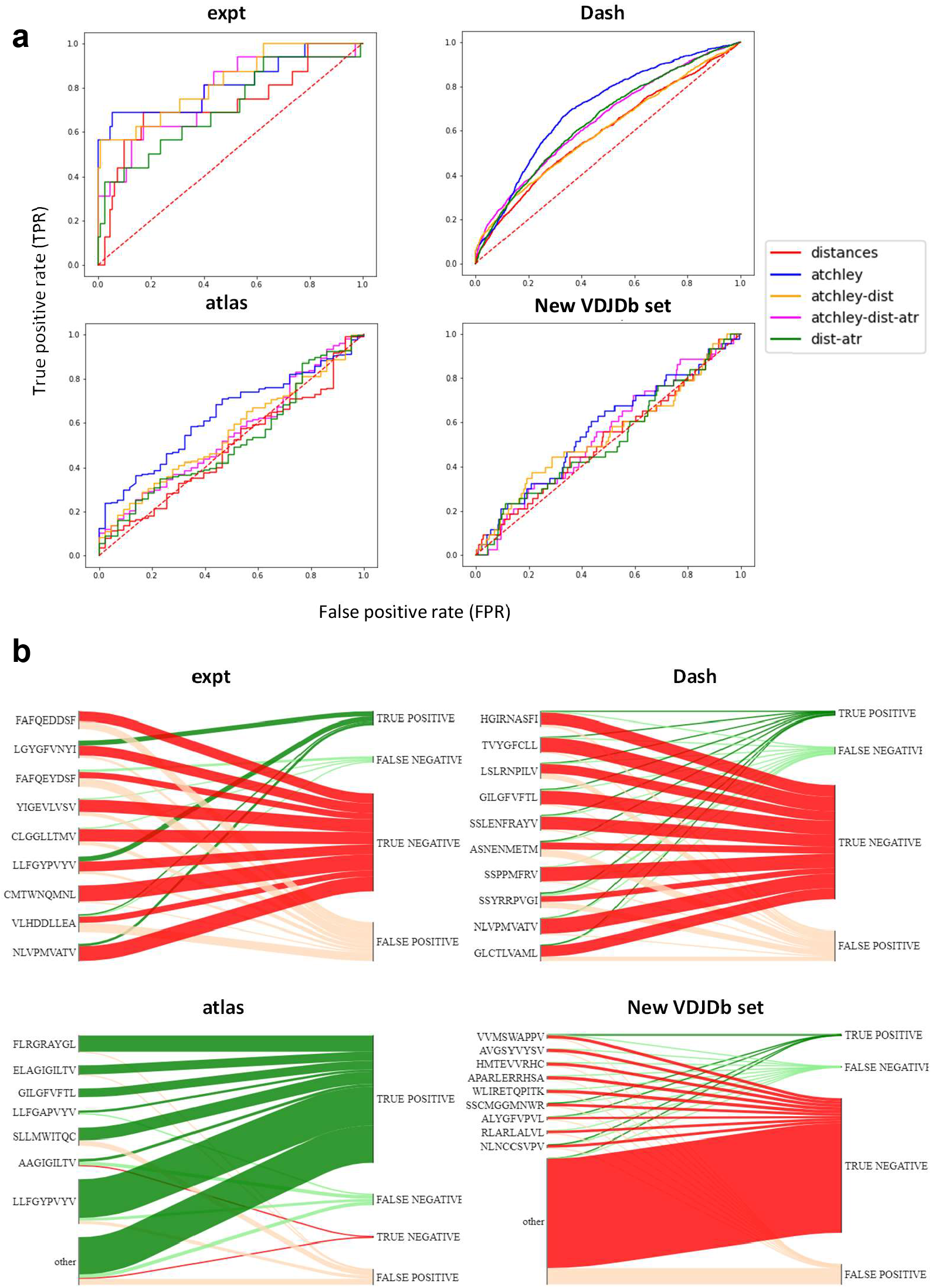
Results of all validation sets used. **a.** ROC curves obtained when the model trained on the STCRDab set are used for prediction on each of the validation sets. **b.** For the model trained on STCRDab using distances only, the diagram shows which proportion of examples from each epitope are classified correctly (true positives and true negatives) or incorrectly (false positives and false negatives) for each of the validation sets used.

**Table S1:** Results of benchmarking on single epitopes. For each epitope, the performance of each tool is calculated (ROC AUC). In each row, the best-performing tool is highlighted in bold and the best-performing model of the ones presented in this paper is boxed.

|  | N pos | N neg | in_pdb | in_vdjb | distances | dist-atr | atchley | atchley-dist | atchley-dist-atr | ImRex | ERGO LSTM | ERGO AE |
| --- | --- | --- | --- | --- | --- | --- | --- | --- | --- | --- | --- | --- |
| VVMSWAPPV | 7 | 120 | no | no | 0.361 | 0.526 | **0.605** | 0.370 | 0.557 | 0.482 | 0.461 | 0.433 |
| ALYGFVPVL | 5 | 122 | no | no | 0.620 | 0.603 | 0.508 | 0.556 | 0.597 | 0.474 | 0.290 | **0.657** |
| HMTEVVRHC | 4 | 123 | no | no | 0.390 | 0.551 | 0.654 | 0.549 | 0.573 | **0.679** | 0.551 | 0.498 |
| APARLERRHSA | 3 | 124 | no | no | 0.559 | 0.570 | 0.449 | 0.538 | 0.495 | **0.901** | 0.591 | 0.562 |
| RLARLALVL | 5 | 122 | no | no | 0.218 | 0.285 | 0.443 | 0.215 | 0.259 | 0.433 | 0.575 | **0.582** |
| NLNCCSVPV | 4 | 123 | no | no | 0.715 | 0.638 | 0.447 | **0.720** | 0.667 | 0.547 | 0.677 | 0.567 |
| RLRAEAQVK | 57 | 336 | no | yes | 0.465 | 0.416 | 0.525 | 0.461 | 0.429 | 0.538 | **0.753** | 0.727 |
| SSPPMFRV | 20 | 1795 | no | yes | 0.393 | 0.412 | 0.396 | 0.373 | 0.424 | 0.665 | **0.891** | 0.814 |
| MLDLQPETT | 6 | 6 | no | yes | 0.750 | 0.333 | 0.306 | 0.417 | 0.250 | **0.778** | 0.694 | 0.583 |
| FLASKIGRLV | 3 | 24 | no | yes | 0.500 | 0.639 | 0.389 | 0.375 | 0.653 | 0.542 | **1.000** | 0.597 |
| TVYGFCLL | 46 | 1839 | no | yes | 0.407 | 0.419 | 0.323 | 0.288 | 0.386 | 0.453 | **0.915** | 0.757 |
| KTWGQYWQV | 3 | 10 | no | yes | 0.800 | 0.633 | 0.700 | 0.867 | 0.633 | 0.433 | **1.000** | 0.933 |
| KLGGALQAK | 324 | 2161 | no | yes | 0.493 | 0.479 | 0.527 | 0.498 | 0.493 | 0.511 | **0.739** | 0.630 |
| AYAQKIFKI | 4 | 62 | no | yes | 0.750 | 0.379 | 0.464 | 0.685 | 0.339 | 0.266 | 0.690 | **0.867** |
| LLDFVRFMGV | 10 | 18 | no | yes | 0.794 | 0.639 | 0.328 | 0.633 | 0.494 | 0.294 | 0.767 | **0.800** |
| HGIRNASFI | 140 | 1674 | no | yes | 0.498 | 0.652 | 0.500 | 0.482 | 0.608 | 0.610 | **0.926** | 0.918 |
| LSLRNPILV | 64 | 1796 | no | yes | 0.437 | 0.443 | 0.644 | 0.456 | 0.465 | 0.520 | **0.902** | 0.745 |
| IVTDFSVIK | 207 | 421 | no | yes | 0.540 | 0.613 | 0.662 | 0.632 | 0.649 | 0.668 | **0.821** | 0.795 |
| RMFPNAPYL | 4 | 12 | no | yes | 0.542 | 0.542 | 0.604 | 0.625 | 0.667 | 0.542 | **0.958** | 0.750 |
| SSYRRPVGI | 455 | 1389 | no | yes | 0.432 | 0.471 | 0.561 | 0.499 | 0.466 | 0.282 | **0.938** | 0.927 |
| AVFDRKSDAK | 175 | 869 | no | yes | 0.465 | 0.441 | 0.494 | 0.460 | 0.432 | 0.534 | **0.716** | 0.669 |
| SLFNTVATLY | 5 | 34 | no | yes | 0.241 | 0.435 | 0.300 | 0.353 | 0.506 | 0.435 | **0.771** | 0.712 |
| RAKFKQLL | 77 | 169 | no | yes | 0.635 | 0.511 | 0.594 | 0.637 | 0.511 | 0.554 | 0.725 | **0.726** |
| FLYALALLL | 7 | 9 | no | yes | 0.508 | 0.635 | 0.190 | 0.444 | 0.349 | 0.349 | **1.000** | 0.968 |
| LGYGFVNYI | 4 | 10 | yes | yes | 0.925 | 0.850 | **1.000** | **1.000** | 0.950 | 0.925 | **1.000** | 0.925 |
| GLCTLVAML | 98 | 1848 | yes | yes | 0.722 | 0.717 | 0.747 | 0.740 | 0.737 | 0.756 | **0.991** | 0.980 |
| LLFGYPVYV | 91 | 36 | yes | yes | 0.865 | 0.867 | 0.888 | 0.888 | 0.876 | 0.908 | **0.935** | 0.922 |
| SLLMWITQC | 33 | 11 | yes | yes | 0.355 | 0.088 | 0.598 | 0.438 | 0.176 | 0.665 | 0.638 | **0.806** |
| SSLENFRAYV | 147 | 1614 | yes | yes | 0.542 | 0.586 | 0.563 | 0.523 | 0.543 | 0.630 | **0.836** | 0.730 |
| GILGFVFTL | 534 | 2028 | yes | yes | 0.722 | 0.741 | 0.841 | 0.779 | 0.785 | 0.822 | **0.982** | 0.969 |
| ELAGIGILTV | 178 | 348 | yes | yes | 0.736 | 0.726 | 0.825 | 0.778 | 0.747 | 0.574 | **0.862** | 0.754 |
| ASNENMETM | 161 | 1717 | yes | yes | 0.518 | 0.609 | 0.468 | 0.461 | 0.608 | 0.486 | **0.948** | 0.900 |
| NLVPMVATV | 63 | 1876 | yes | yes | 0.623 | 0.648 | 0.558 | 0.628 | 0.626 | 0.495 | **0.987** | 0.956 |

